# The molecular chaperone DnaK accelerates protein evolution

**DOI:** 10.1101/040600

**Authors:** José Aguilar-Rodríguez, Beatriz Sabater-Muñoz, Víctor Berlanga, David Alvarez-Ponce, Andreas Wagner, Mario A. Fares

**Affiliations:** Institute of Evolutionary Biology and Environmental Studies, University of Zurich, Zurich, Switzerland; Swiss Institute of Bioinformatics, Lausanne, Switzerland; Department of Genetics, Smurfit Institute of Genetics, University of Dublin, Trinity College Dublin, Dublin, Ireland; Instituto de Biología Molecular y Celular de Plantas (CSC-UPV), Valencia, Spain; Department of Biology, University of Nevada, Reno, USA; Santa Fe Institute, Santa Fe, New Mexico, USA

**Keywords:** DnaK, Hsp70 chaperones, molecular chaperones, mutational robustness, experimental evolution, protein evolution, *Escherichia coli*, evolutionary rates

## Abstract

Molecular chaperones, also known as heat-shock proteins, refold misfolded proteins and help other proteins reach their native conformation. Thanks to these abilities, some chaperones, such as the Hsp90 protein or the chaperonin GroEL, can buffer the deleterious phenotypic effects of mutations that alter protein structure and function. Hsp70 chaperones use a chaperoning mechanism different from Hsp90 and GroEL, and it is not known whether they can also buffer mutations. Here, we show that they can. To this end, we performed a mutation accumulation experiment in *Escherichia coli*, followed by whole-genome resequencing. Our sequence data shows that overexpression of the Hsp70 chaperone DnaK increases the tolerance of its clients for nonsynonymous nucleotide substitutions and nucleotide insertions and deletions. We also show that this elevated mutational buffering on short evolutionary time scales translates into differences in evolutionary rates on intermediate and long evolutionary time scales. To this end, we compared the evolutionary rates of DnaK clients and nonclients using the genomes of *E. coli*, *Salmonella typhimurium*, and 83 other gamma-proteobacterial species. We find that clients that interact strongly with DnaK evolve faster than weakly interacting clients. Our results imply that all three major chaperone classes can buffer mutations and affect protein evolution. They illustrate how an individual protein like a chaperone can have a disproportionate effect on proteome evolution.

## Introduction

Molecular chaperones assist proteins in reaching their native conformations, prevent protein aggregation, and refold misfolded proteins (Young et al. 2004; Hartl and Hayer-Hartl 2009; Hartl et al. 2011). Thanks to these roles, chaperones can restore the native conformation of proteins destabilized by environmental perturbations, thus providing environmental robustness to organisms coping with stressful conditions. Because some chaperones can buffer the deleterious effects of mutations that affect protein folding, they are also a source of mutational robustness (Bogumil and Dagan 2012; Fares 2015).

There are three main chaperone systems, which are the Hsp90 system, the Hsp70 system, and the Hsp60 system (or chaperonins), of which the bacterial GroEL is a prominent member (Hartl et al. 2011). Overwhelming evidence shows that Hsp90 and GroEL can buffer mutations (Bogumil and Dagan 2012), but whether the same holds for any major chaperone from the Hsp70 system is to our knowledge unknown.

Pioneering work carried out by Rutherford and Lindquist (1998) showed that inhibition of the chaperone Hsp90 can unveil cryptic genetic variation ‒ genotypic variation without phenotypic variation ‒ in the fruit fly *Drosophila melanogastert*. Subsequently, similar observations have been made in the plant *Arabidopsis thaliana* (Queitsch et al. 2002), the yeast *Saccharomyces cerevisae* (Cowen and Lindquist 2005) and the fish *Astyanax mexicanus* (Rohner et al. 2013). Further support was recently provided by Burga et al. (2011) who found that high induction of Hsp90 during development of the nematode *Caenorhabditis elegans* reduced the penetrance of certain mutations. Additionally, Lachowiec et al. (2013) found that paralogs of duplicated kinase-coding genes that encode a substrate of Hsp90 (i.e., a Hsp90 “client”) in *A. thaliana* often evolve faster than nonclient paralogs. In general, the rate at which non-conservative substitutions ‒ those that alter physicochemical properties of amino acids ‒ accumulate is especially accelerated in Hsp90 clients (Pechmann and Frydman 2014).

Multiple studies also demonstrate mutational buffering mediated by the bacterial chaperonin GroEL. For example, Fares et al. (2002) showed that overexpressing GroEL considerably improved the fitness of *Escherichia coli* strains with a high load of deleterious mutations, a pattern that was also observed later in *Salmonella typhimurium* (Maisnier-Patin et al. 2005). Moreover, GroEL overexpression in *E. coli* increases the ability of GroEL client proteins to tolerate mutations (Tokuriki and Tawfik 2009; Bershtein et al. 2013; Wyganowski et al. 2013; Sabater-Muñoz et al. 2015), as well as their ability to undergo adaptive evolution (Tokuriki and Tawfik 2009; Wyganowski et al. 2013). Buffering of destabilizing mutations accelerates the evolutionary rates of GroEL clients (Bogumil and Dagan 2010; Warnecke and Hurst 2010; Williams and Fares 2010; Pechmann and Frydman 2014).

While no Hsp70 chaperone has been directly implicated in mutational buffering, pertinent circumstantial evidence exists. For example, DnaK ‒ the major bacterial Hsp70 chaperone ‒ is overexpressed together with GroEL in *S. typhimurium* lineages with deteriorated fitness caused by accumulating deleterious mutations (Maisnier-Patin et al. 2005). In addition, *D. melanogaster* populations showing inbreeding depression, where increased homozygosity exposes recessive deleterious mutations, significantly up-regulate the expression of Hsp70 compared to outbred populations (Pedersen et al. 2005).

The chaperones from the Hsp70 system are very conserved from bacteria to humans (Powers and Balch 2013). They play a central role in proteome integrity, and are involved both in co- and post-translational folding (Hartl et al. 2011). In bacteria, the Hsp70 chaperone DnaK (together with GroEL and the Trigger Factor) is one of the main molecular chaperones, where it is the central hub in the chaperone network of the cytosol (Bukau and Walker 1989; Calloni et al. 2012). It interacts with ~700 mostly cytosolic proteins (Calloni et al. 2012). DnaK is highly expressed constitutively, but also stress inducible, and essential at 42°C (Bukau and Walker 1989; Calloni et al. 2012). During its ATP-dependent reaction cycle, DnaK interacts with the Hsp40 co-chaperone DnaJ, and the nucleotide exchange factor GrpE (Hartl et al. 2011). DnaJ determines the client binding specificity of DnaK (Straus et al. 1990; Hoffmann et al. 1992).

Most mutations affecting proteins are neutral or deleterious (Eyre-Walker and Keightley 2007), and functionally important mutations often destabilize proteins (Tokuriki et al. 2008; Wyganowski et al. 2013). If DnaK buffers destabilizing mutations, then the deleterious effects of mutations in highly interacting (strong) clients should be lower than in sporadic (weak) clients, where they should be lower than in nonclients. In other words, the higher the dependency of a protein’s integrity on DnaK, the higher should be its tolerance to mutations, and the lower the signature of purifying selection that purges those mutations. With this reasoning in mind, we here use laboratory experiments to evaluate the effect of DnaK buffering on the evolution of its client proteome on short evolutionary time scales. We complement our experimental observations with sequence analyses to study the effect of DnaK on intermediate and long evolutionary time scales.

## Results

### Experimental evolution of E. coli under DnaK overexpression

To study the effect of DnaK overexpression on protein evolution experimentally, we performed mutation accumulation experiments similar to those we reported recently for the chaperonin GroEL, but for DnaK overexpression (Sabater-Muñoz et al. 2015). Briefly, we initiated six parallel and independent clonal lines of evolution, all of which derived from the same hypermutable clone (*E. coli* K12 MG1655 ∆*mutS*) (Bjedov et al. 2007). Cells of the six lines all harbored the plasmid pKJE7, which contains the operon *dnaK-dnaJ-grpE* under the control of the L-arabinose-inducible *araB* promoter (Nishihara et al. 1998). We evolved all six lines through repeated single-cell bottlenecks in the presence of the inducer, to ensure overexpression of DnaK (DnaK^+^ lines), as well as of the cochaperone DnaJ and the nucleotide exchange factor GrpE. Because of the bottlenecks we exposed the populations to, genetic drift was strong and the efficiency of selection was weak during the experiment, such that non-lethal mutations are free to accumulate (Barrick and Lenski 2013). We evolved three of the six clonal lines at 37°C, and the other three at 42°C. The higher temperature serves to increase the deleterious effect of destabilizing mutations in the bacterial proteome (Bukau and Walker 1989). At each of the two temperatures, we also evolved a control clonal line founded from the same parental strain, but carrying a pKJE7-derived plasmid where the operon *dnaK-dnaJ-grpE* is deleted. The two control lines therefore cannot overexpress DnaK, even though their growth medium contains L-arabinose (DnaK^−^ lines).

The overexpression of DnaK may be energetically costly, just as is the case for the chaperonin GroEL (Fares et al. 2002; Sabater-Muñoz et al. 2015). In principle, this cost could favor the accumulation of mutations that lead to a decrease in the expression of DnaK during the evolution experiment. However, we observed that the overexpression of DnaK was maintained through the mutation accumulation experiment at both 37°C and 42°C (Fig. 1, Supplementary Fig. 1). In the presence of the inducer L-arabinose, all DnaK^+^ lines overexpressed DnaK not only at the start of the evolution experiment, but also at the end, except for a DnaK^+^ line evolved at 42°C. However, this loss of overexpression occurred towards the end of the experiment and even then there was still overexpression for most of the daily growth cycle of this line (Supplementary Fig. 2). In no line did we observe overexpression in the absence of the inducer. The control DnaK^-^ lines always exhibited wild-type expression levels of DnaK.

**Figure 1.**
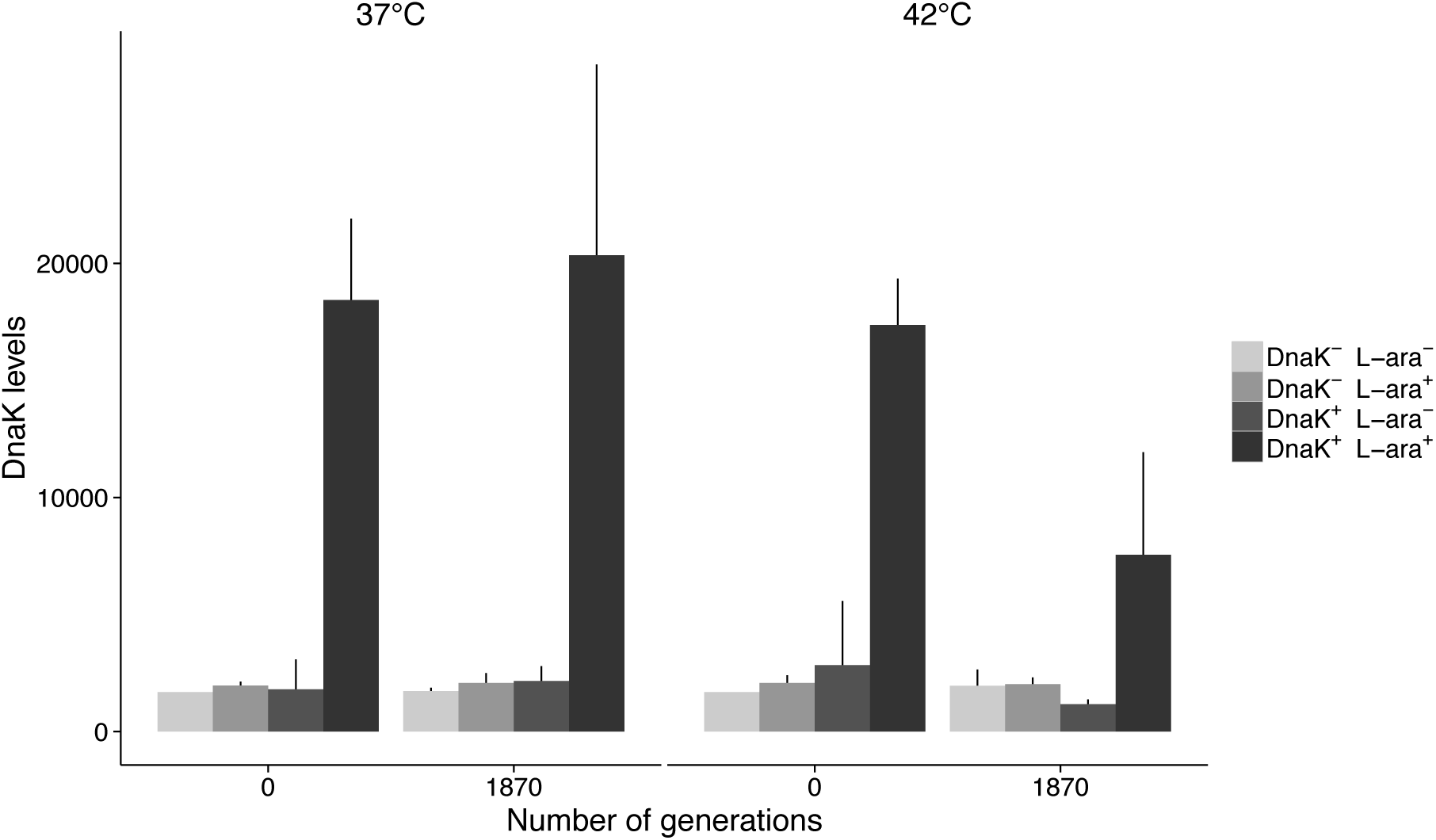
DnaK abundance at the end of the mutation accumulation experiment. We measured the abundance of the chaperone DnaK for both DnaK^+^ and DnaK^−^ lines evolved for approximately 1,870 generations at 37°C and 42°C. For comparison, we also measured the abundance of the chaperone in the ancestral DnaK^+^ and DnaK^−^ strains at both temperatures. We determined DnaK levels in the presence and absence of the inducer L-arabinose (L-ara^+^ and L-ara^−^, respectively), as described in Material and Methods (“Verification of DnaK overexpression”), via the intensity of the DnaK band in a Western blot. The evolved lines did not lose the ability to overexpress DnaK in the presence of the inducer L-arabinose except for a DnaK^+^ line evolved at 42°C (line #2), which explains the decrease in the average DnaK abundance at the end of the evolution experiment. However, this loss of overexpression occurred late in the evolution experiment, and it is not even complete for most of the daily growth cycle of this line (Supplementary Figure 2). The height of the bars indicates mean DnaK abundance across two experimental replicates per strain and condition. Error bars represent one standard deviation of the mean.

Importantly, while all DnaK^-^ lines underwent repeated extinctions shortly after 85 single-cell bottlenecks or approximately 1,870 generations of evolution, the DnaK^+^ lines overexpressing DnaK could be passaged for as long as 180 daily passages or ~3,960 generations. This observation suggests that overexpressing the chaperone DnaK has increased the robustness of the cells to the accumulation of deleterious mutations.

### Overexpressing DnaK increases the robustness to protein-changing mutations of DnaK clients

Together with the ancestral genome, we sequenced the genomes of all eight clonal lineages after ~1,870 generations. In the genomes of the evolved DnaK^+^ lines, we first studied the incidence of protein-changing mutations ‒ those that include nonsynonymous nucleotide substitutions as well as nucleotide insertions and deletions (indels) ‒ among DnaK clients and nonclients. In addition to nonsynonymous substitutions, we also analyzed indels, since this mutation type, which can affect protein structure and folding, could in principle be buffered by a molecular chaperone such as DnaK. In this analysis, we considered as nonclients all proteins from the *E. coli* proteome that are not part of the set of 671 DnaK clients recently determined by Calloni et al. (2012), and analyzed the lines evolved at 37°C and 42°C independently. To improve statistical power, we combined mutations across DnaK^+^ lines evolved at the same temperature after classifying them according to whether they affect DnaK clients or nonclients. In total, we found that the lines evolved at 37°C harbored 74 protein-changing mutations in clients and 324 mutations in nonclients (Supplementary Table 1). In other words, 11.03% of clients and 9.73% of nonclients were mutated. At this temperature, although the proportion of mutated clients is greater than the proportion of mutated nonclients, this difference is not significant (Fisher’s exact test: odds ratio *F* = 1.15, *P* = 0.32). The DnaK^+^ lines evolved at 42°C accumulated 82 mutations in clients and 387 in nonclients (Supplementary Table 1). In this case, 12.22% of clients and 11.63% of clients were affected by mutations. Similarly to our observations at 37°C, we found no evidence for a larger proportion of mutated clients (Fisher’s exact test: odds ratio *F* = 1.06, *P* = 0.64). The same held at both temperatures when we controlled for the number of nonsynonymous sites in clients (502,499 sites) and nonclients (2,692,140 sites) (Fisher’s exact test: odds ratio *F* = 1.22, *P* = 0.13 at 37°C and *F* = 1.14, *P* = 0.31 at 42°C).

While these analyses show that the relative number of mutations in clients is higher than in nonclients, the fact that this difference is not significant suggest that DnaK clients in *E. coli* do not have a greater tolerance to protein-changing mutations than nonclients. However, these analyses ignore a potentially important confounding factor: Clients may be intrinsically more sensitive to mutations than nonclients. Indeed, it has been shown that nonclient proteins are more soluble and expose less hydrophobic regions to the solvent than clients, that is client proteins are intrinsically less stable than nonclient proteins (Calloni et al. 2012). If so, comparing clients and nonclients directly would not be suitable to identify the masking of deleterious mutations by DnaK. The reason is that while the proportion of mutated clients is not significantly higher than the proportion of mutated nonclients, this proportion may have increased as a consequence of DnaK overexpression. To evaluate this possibility, we compared the number of mutations among DnaK clients and nonclients in the DnaK^+^ lines to the same numbers in the DnaK^−^ lines (Supplementary Table 1). In the DnaK^−^ line evolved at 37°C, 13% of mutations (19 out of 141) affected DnaK clients. Compared to this expected proportion when DnaK is not overexpressed, the proportion of mutations in clients in the DnaK^+^ lines is significantly higher than expected (74 out of the total 394 mutations, 18.6%; binomial test: *P* = 0.002; Fig. 2). Similarly, compared to the DnaK^−^ line evolved at 42°C, where 11% of all mutations affected DnaK clients (15 out of 136 mutations), the DnaK^+^ lines showed significantly more mutations in clients (84 out of 471 total mutations, 17.8%; binomial test: *P* = 1.19 × 10^-5^; Fig. 2). These results suggest that overexpressing DnaK does indeed increase the robustness of its clients to mutations.

**Figure 2.**
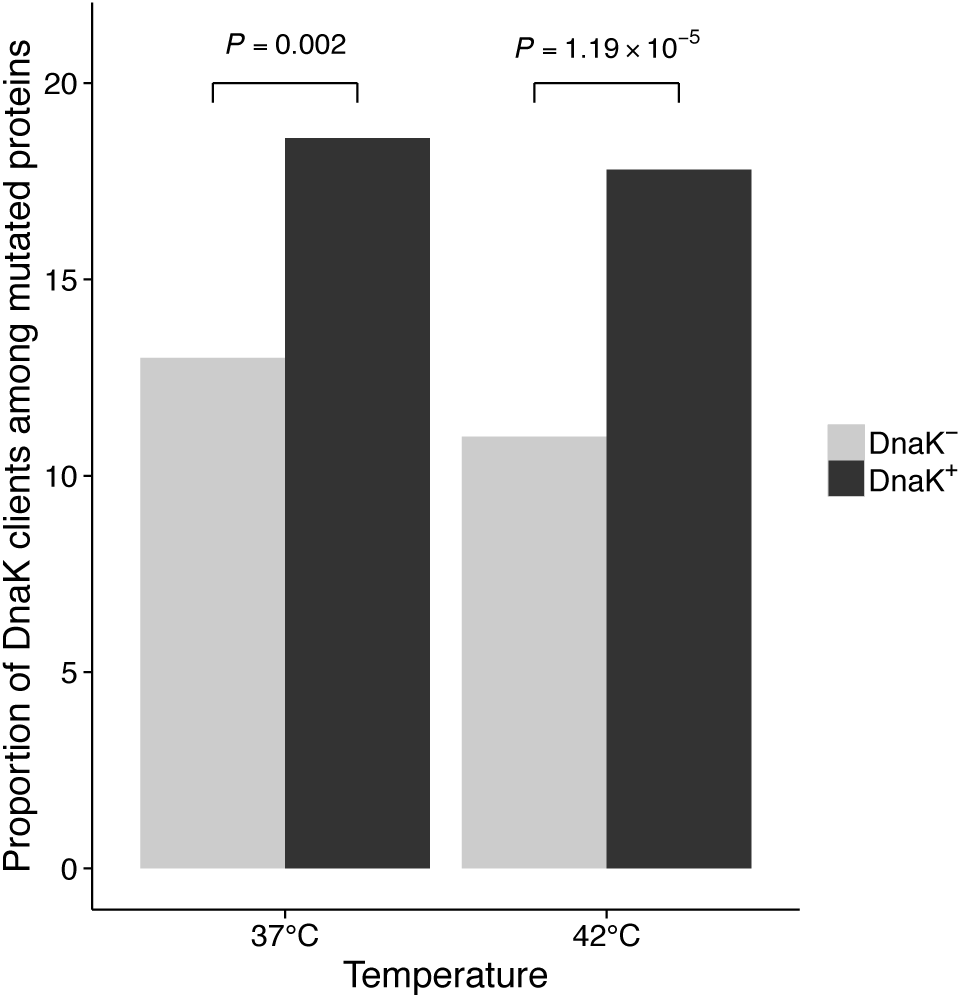
Higher proportion of mutated DnaK clients among all mutated proteins after ~1,870 generations of mutation accumulation. The proportion of mutated DnaK clients is significantly higher in DnaK^+^ lines that overexpress DnaK than in the control DnaK^−^ lines that do not express the chaperone at such high levels. This is observed both for lines evolved at 37°C and 42°C. We consider a protein as mutated if it has a nonsynonymous substitution and/or an indel. Statistical significance was evaluated with a binomial test.

Temperature itself had no significant effect on the proportion of mutated clients and nonclients (Fisher’s exact test: odds ratio *F* = 1.08, *P* = 0.72). At both temperatures, the number of accumulated mutations in strong DnaK clients was approximately twice as high as that in weak clients, but this difference was not significant, likely because of a lack of statistical power (Supplementary Table 1, Supplementary Information section 2.1). Importantly, this observation is not spuriously explained by a difference in coding sequence length between strong and weak clients (Wilcoxon rank-sum test, *P*-value = 0.512).

### DnaK accelerates protein evolution on intermediate and long evolutionary time scales

We wanted to find out if the DnaK-mediated mutational buffering we observed on the short time scales of laboratory evolution has also left signatures on longer evolutionary time scales. To this end, we determined two measures of evolutionary rates for protein-coding genes from gamma-proteobacteria (Supplementary Table 2). The first, nonsynonymous divergence among one-to-one orthologs of *E. coli* and *Salmonella enterica*, is relevant for intermediate evolutionary time scales. The second, protein distance among orthologous proteins found in 85 gamma-proteobacterial genomes (including *E. coli* and *S. enterica*), is relevant for long time scales. We employ protein distance instead of nonsynonymous distance because amino acid replacements are less sensitive than nucleotide substitutions to the expected loss of phylogenetic signal between sequences of distantly related taxa. To assess how strongly a protein depends on DnaK for folding, we used recent experimental proteomic data which determined how strongly 671 DnaK-interacting proteins interact with DnaK by measuring the fraction of cellular protein bound to DnaK at 37°C, a property that correlates with chaperone dependency for folding and maintenance and residence time of the protein on DnaK (Calloni et al. 2012). We note that this interaction strength is more likely to have remained unchanged during the divergence of *E. coli* and *S. enterica*, than during the divergence of all the other 83 gamma-proteobacterial species we analyzed.

We find a moderately strong but highly significant positive association between DnaK dependency and the rate of nonsynonymous substitutions for *S. enterica* and *E.coli* (Spearman’s ρ = 0.367, *N* = 627, *P-*value < 2.2×10^-16^; Fig. 3A). This indicates that the stronger the interaction of a protein with DnaK, the faster the protein evolves. The same pattern is obtained at the larger time scales of protein distances for 85 gamma-proteobacterial genomes (Spearman’s ρ = 0.257, *N* = 311, *P-*value = 4.4×10^-6^; Fig. 3B). Gene expression level, which is the most important determinant of protein evolutionary rates, at least in unicellular organisms (Pál et al. 2001; Drummond et al. 2005), is a possible confounding factor in this analysis. For example, using codon usage bias (CUB) as a proxy for gene expression, we observe that genes with higher CUB show lower nonsynonymous divergence (Spearman’s ρ = −0.558, *N* = 1014, *P-*value < 2.2×10^-16^), protein distance (Spearman’s ρ = −0.255, *N* = 3159, *P-*value < 2.2×10^-16^) and DnaK dependency (Spearman’s ρ = −0.262, *N* = 627, *P-*value = 2.5×10^-11^). However, the association between DnaK dependency and evolutionary rate cannot be solely explained by this confounding factor: A partial correlation analysis shows that the association still holds after controlling for CUB, both on intermediate time scales (Spearman’s ρ = 0.295, *N* = 627, *P-*value = 1.2×10^-14^) and long time scales (Spearman’s ρ = 0.229, *N* = 311, *P-*value = 3.8×10^-5^). We use CUB instead of gene expression data here for two main reasons. First, we can compute CUB for 627 DnaK clients, while expression data is only available for 457 clients. Second, gene expression data has been measured in just one environment and one strain of *E. coli*, while CUB is the result of selective pressures imposed by many different environments over long periods of time. However, the association between evolutionary rate and DnaK dependency still holds after correcting for gene expression directly (Supplementary Information section 2.2). Together, these results indicate that the chaperone DnaK affects protein evolution in accordance with the mutational buffering hypothesis. Importantly, this effect is not only independent of CUB and gene expression, but also of other biological factors, such as essentiality and number of protein-protein interactions (Supplementary Information section 2.3, Supplementary Table 3).

**Figure 3.**
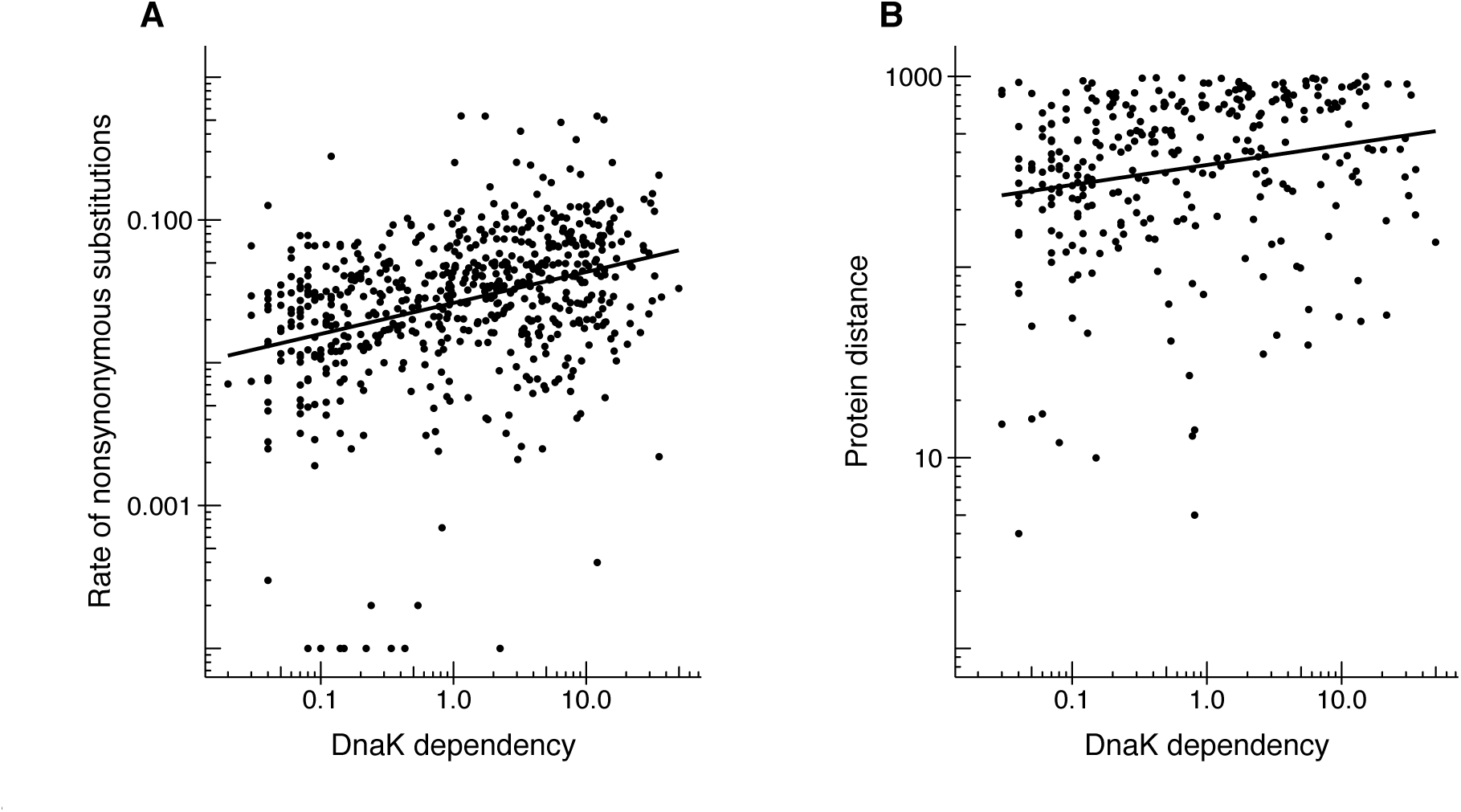
DnaK accelerates protein evolution on intermediate and long evolutionary time scales. Scatter-plots showing the relationship between DnaK dependency (calculated as a relative enrichment factor that indicates the fraction of cellular protein bound to DnaK at 37°C, horizontal axis) and the degree of divergence over (**A**) intermediate time scales, measured as nonsynonymous divergence (Spearman’s ρ = 0.367, *N* = 627, *P-*value < 2.2×10^-16^), and (**B**) long time scales, measured as protein distance (Spearman’s ρ = 0.257, *N* = 311, *P-*value = 4.4×10^-6^) (vertical axes). Solid lines represent the best fit to the points. Note the logarithmic scale on both axes.

Finally, while we observe that strong clients evolve faster than weak clients on both evolutionary time scales, we find that clients evolve more slowly than nonclients (Supplementary Information section 2.4, Supplementary Fig. 3). This last difference cannot be explained by the number of protein-protein interactions, by essentiality, or by CUB as confounding factors (Supplementary Information section 2.4, Supplementary Tables 4 and 5). The reason for this observation could be that clients are intrinsically less robust to mutations than nonclients due to some general physicochemical difference. For example, Calloni et al. (2012) found that DnaK clients have generally low solubility, often belong to heterooligomeric complexes, and are prone to misfolding.

## Discussion

We show how the constant overexpression of the DnaK-DnaJ-GrpE chaperone system over the course of a mutation accumulation experiment increases the number of protein-changing mutations ‒ nonsynonymous substitutions and indels ‒ affecting DnaK clients. Additional evidence of mutational buffering by DnaK is provided by the observation that only evolving lines overproducing this chaperone avoid extinction after experiencing more than 85 single-cell bottlenecks. Recently, we obtained similar results in hypermutable *E. coli* cells evolving in identical conditions but overproducing the GroEL-GroES chaperonin system (Sabater-Muñoz et al. 2015). There, we observed that lines evolving with high levels of GroEL were not only less prone to extinction under strong genetic drift than control lines, but also that they were accumulating significantly more indels and replacements between amino acids belonging to different physicochemical categories.

Additionally, we find that DnaK-mediated mutational buffering has left a trace in DnaK clients during the divergence of 85 different gamma-proteobacterial species over much longer evolutionary time scales than those explored in our laboratory evolution experiment. We find that clients that depend more on DnaK for folding tend to evolve faster than less interacting clients. Similar chaperone-mediated accelerations of protein evolution have been observed on GroEL clients (Bogumil and Dagan 2010; Warnecke and Hurst 2010; Williams and Fares 2010; Bogumil and Dagan 2012) and Hsp90 clients (Lachowiec et al. 2013; Pechmann and Frydman 2014). However, we notice that DnaK clients evolve slower than nonclients. This is likely the result of important physicochemical differences between clients and nonclients. For example, clients are prone to aggregation and misfolding (Calloni et al. 2012), which may make them intrinsically less robust to destabilizing mutations.

Despite the great differences in the mechanism of chaperone action between the three major chaperone families – chaperonins, Hsp90 chaperones and Hsp70 chaperones – (Young et al. 2004; Hartl et al. 2011; Bogumil and Dagan 2012; Kim et al. 2013), at least some of their members seem to have qualitatively comparable effects on protein evolution. Elucidating to what extent the buffering mechanisms of these chaperones differ is an important future direction of enquiry.

Thanks to their fostering of mutational robustness, chaperones can facilitate evolutionary innovations (Rutherford 2003), even though we do not study such innovations here. The increase in the mutational robustness of a protein caused by chaperone interactions reduces the efficiency of purifying selection in purging mutations in the protein. Thanks to chaperone-mediated buffering, many such mutations are neutral and can persist in a population. Importantly, these cryptic genetic variants may include preadaptive mutations that can generate evolutionary innovations in new environments (Tokuriki and Tawfik 2009; Wyganowski et al. 2013). To illuminate if and how DnaK can increase the innovability of its client proteome will also be an interesting subject for future work.

In summary, we analysed evolutionary rates of proteins that are subject to DnaK-assisted folding on short, intermediate, and long evolutionary time scales through a combination of experimental and comparative approaches. Most of our evidence indicates that the bacterial chaperone DnaK can buffer mutations in its client proteins, and that these proteins therefore evolve faster than in the absence of DnaK-mediated folding. This is, to our knowledge, the first demonstration that a member of the Hsp70 family can buffer the effect of mutations, with long-term consequences on protein evolution (Bogumil and Dagan 2012). Through its role in protein folding, an individual chaperone such as DnaK can have a disproportionate effect on proteome evolution, and thus on genome evolution.

## Materials and Methods

### Sequence data

We obtained the complete genomes of *E. coli* K-12 MG1655 (NC_000913) and *S. enterica* serovar Typhimurium LT2 (NC_003197) from GenBank Genomes (ftp://ftp.ncbi.nih.gov/genomes/Bacteria/). We also used a dataset from Williams and Fares (2010) that consists of 1092 multiple sequence alignments of conserved orthologous proteins from 85 gamma-proteobacterial genomes.

### Orthology

We identified 3159 one-to-one orthologs in *E. coli* and *S. enterica* genomes as reciprocal best hits (Tatusov et al. 1997) using the Basic Local Alignment Search Tool (BLAST, i.e., BLASTP with an E-value cut-off of 10^-10^). We aligned each pair of orthologous proteins with the Needleman-Wunsch dynamic programming algorithm, using the Needle program from the EMBOSS package (Rice et al. 2000). We translated the resulting alignments into codon-based nucleotide alignments with PAL2NAL (Suyama et al. 2006).

### Evolutionary rates

We estimated the rate of nonsynonymous substitutions (*d*_N_) using the program codeml from the package PAML 4.7 (one-ratio model M0) (Yang 2007). We calculated protein distances for the gamma-proteobacterial alignments from Williams and Fares (2010), using PROTDIST from the PHYLIP package (Felsenstein 2005) and the Jones, Taylor and Thornton (JTT) substitution matrix (Jones et al. 1992). We calculated an average distance for each cluster of orthologous proteins as the mean of all pairwise distances.

### Codon usage bias

We computed the Codon Adaptation Index (CAI) using the program CAI from the EMBOSS package (Rice et al. 2000). We calculated Codon Usage Bias (CUB) for each pair of *E. coli* - *S. enterica* orthologs as the mean of the CAI values for each pair of orthologs. We used CUB as a proxy for gene expression.

### DnaK dependency

We obtained information about DnaK clients from Calloni et al. (2012). This study used quantitative proteomics to identify 671 DnaK interactors or client proteins. For each of these proteins, the investigators calculated a relative enrichment factor that indicates the fraction of cellular protein bound to DnaK at 37°C. We used this measure as a proxy for DnaK dependency.

### Bacterial strains and plasmids

We obtained *E. coli* K-12 substr. MG1655 ∆ *mutS* from Ivan Matic (Université Paris Descartes, INSERM U1001, Paris, France) through Jesús Blázquez (Centro Nacional de Biotecnología, CSIC, Madrid, Spain) (Bjedov et al. 2007). In this *E. coli* strain the gene encoding the protein MutS has been deleted. This protein is a component of the mismatch repair system that recognizes and binds mispaired nucleotides so that the mispairing can be corrected by two further repair proteins, MutL and MutH. The strain MG1655 ∆*mutS* has a predicted mutation rate that is 1000-fold higher than the wild type (Bjedov et al. 2007; Turrientes et al. 2013), which ensures that a sufficient number of mutations occur during the mutation accumulation experiment. We transformed this strain with the plasmid pKJE7 (Takara, Cat. #3340), which contains an operon encoding DnaK, and its co-chaperones DnaJ and GrpE under the regulation of a single promoter inducible by L-arabinose (Nishihara et al. 1998). We generated a control strain by transforming the same ∆*mutS* strain with a plasmid that lacks the operon *dnaK*-*dnaJ*-*grpE* but is otherwise identical to pKJE7. We refer to this plasmid as pKJE7-DEL(*dnaK*-*dnaJ*-*grpE*). This control plasmid was derived from the plasmid pKJE7 by removal of the operon *dna*K-*dna*J-*grp*E with a restriction digest using *Bam*HI and *Spe*I, followed by religation, after obtaining permission for plasmid modification from Takara.

### Evolution experiment

We evolved six clonal lines of the hypermutable *E. coli* ∆*mutS* strain containing pKJE7 (DnaK^+^ lines) and two lines containing the control plasmid pKJE7-DEL(*dnaK*-*dnaJ*-*grpE*) (DnaK^-^ lines) by daily passaging them through single-cell bottlenecks on solid LB medium (agar plates; Pronadisa #1551 and #1800) supplemented with 20 μg/mL of chloramphenicol (Sigma-Aldrich #C0378) and 0.2% (w/v) of L-arabinose (Sigma-Aldrich #A3256). We passaged both the DnaK^-^ and DnaK^+^ lines during 85 days or approximately 1,870 generations (assuming ~22 generations per daily growth cycle). We evolved half of the DnaK^+^ and DnaK^-^ lines under mild heat-stress (42°C) while the other half remained at 37°C.

### Verification of DnaK overexpression

We grew the ancestral and evolved strains (DnaK^+^ and DnaK^-^, at 37°C and 42°C) from glycerol stocks in liquid LB medium supplemented with 20 μg/mL of chloramphenicol in the presence or absence of the inducer L-arabinose (0.2%). After 24h of growth, we pelleted cells by centrifugation at 12000 rpm. We resuspended the pelleted cells in 100μl lysis buffer (containing 200 mM Tris-HCl pH6.8, 10mM DTT, 5% SDS, 50% glycerol). To prepare a crude extract, we first boiled resuspended cells at 95°C for 15 minutes. After the removal of cell debris by centrifugation, we quantified soluble proteins using the Bradford method (Bradford 1976). We loaded one microgram of total protein for each sample in SDS-PAGE gels (12.5% resolving gel). In addition, we loaded onto all gels samples from the ancestral DnaK^-^ and DnaK^+^ strains grown in the presence of inducer at 37°C, as controls to facilitate inter-gel comparisons. We detected DnaK protein by Western blotting using as primary antibody a mouse monoclonal antibody specific to *E. coli* DnaK (Abcam #ab69617) at a 1:10,000 dilution, and as secondary antibody a goat polyclonal (alkaline phosphatase-conjugated) antibody specific to mouse IgG1 (Abcam #ab97237). We scanned membranes after colorimetric detection of conjugated antibodies with the BCIP^®^/NBT-Purple liquid substrate system (Sigma-Aldrich #BP3679), and used ImageJ to quantify the intensity of DnaK bands on the Western blots (Schneider et al. 2012). We used the control samples to normalize abundances, which allow the comparison of DnaK levels across experiments.

We examined the change in DnaK levels along a daily cycle of growth for a DnaK^+^ line evolved at 42°C (line #2, Supplementary Figure 2) that lost overexpression after ~1,870 generations of mutation accumulation. After 24h of exponential growth at 42°C in liquid LB medium supplemented with chloramphenicol, we diluted the culture to OD ~0.3, and induced DnaK expression by adding 10 mM of L-arabinose. We allowed the culture to grow for another 24h in the presence of this expression inducer. Each hour we removed one mililiter of culture and measured the DnaK level following the protocol described above.

### Whole-genome resequencing

We sequenced the genomes of the DnaK^-^ and DnaK^+^ lines after 85 single-cell bottlenecks, an addition to the ancestral ∆*mutS* strain from which both the DnaK^+^ and DnaK^-^ lines were derived.

Specifically, for the evolved lines we performed paired-end Illumina whole-genome sequencing. For DNA extraction, we used the QIAmp DNA mini kit (Qiagen, Venlo [Pays Bas], Germany) in a QiaCube automatic DNA extractor using bacterial pellets obtained from approximately 10 mL cultures. We constructed multiplexed DNAseq libraries from each clonal evolution line using the TrueSeq DNA polymerase chain reaction-free HT sample preparation kit (Illumina). We performed paired-end sequencing on an Illumina HiSeq2000 platform, using a 2 × 100 cycles configuration.

We performed whole-genome sequencing of the ancestral strain using paired-end and shotgun 454 sequencing (Roche/454 Life Sciences, Branford, CT). We performed a *de novo* assembly using the GS De Novo Assembler version 2.6 (Roche), and combined the mapped and *de novo* assemblies into a single assembly using in-house scripts. We examined reads to confirm indels using ssaha and breseq v0.24rc4 (Deatherage and Barrick 2014), and transferred annotations from *E. coli* MG1655 using RATT (Otto et al. 2011).

We converted sequencing reads converted from Illumina quality scores into Sanger quality scores. Subsequently, we used the breseq v 0.24rc4 (version 4) pipeline (Deatherage and Barrick 2014) for aligning the Illumina reads to our *E. coli* parental genome and for identifying single nucleotide polymorphisms and indels using bowtie2 (Langmead and Salzberg 2012). We performed individual runs of breseq, with junction prediction disabled but otherwise default parameters, for the ancestral sequence, as well as for each of the evolved lines.

## Acknowledgments

The authors thank Xiaoshu Chen and Jianzhi Zhang for kindly providing us with the gene expression data. This work was supported by the Forschungskredit program of the University of Zurich (grant FK-14-076 to J.A.), the Swiss National Science Foundation (grant 31003A_146137 to A.W.), the University Priority Research Program in Evolutionary Biology at the University of Zurich (to A.W.), the Science Foundation Ireland (grant 12/IP/1673 to M.A.F), and the Spanish Ministerio de Economía y Competitividad (grant BFU2012-36346 to M.A.F.).

